# Selection of lamb size and early pregnancy in Soay sheep (*Ovies aries*)

**DOI:** 10.1101/2020.09.16.299685

**Authors:** Maria João Janeiro, Jonathan M. Henshaw, Josephine M. Pemberton, Jill G. Pilkington, Michael B. Morrissey

## Abstract

The paradox of stasis – the unexpectedly slow evolution of heritable traits under direct selection – has been widely documented in the last few decades. This paradox is often particularly acute for body size, which is often heritable and where positive associations of size and fitness are frequently identified, but constraints to the evolution of larger body sizes are often not obvious. Here, we identify a trade-off between survival and size-dependent reproduction in Soay sheep (*Ovis aries*), contributes to selection against large body size. Using recently developed theory on non-linear developmental systems, then decompose total selection of ewe lamb mass along different causal paths to fitness. Larger lambs are more likely to become pregnant, which has a large viability cost. After controlling for this pathway, however, the association between lamb mass and subsequent lifetime fitness is positive. Thus this trade-off does not fully explain stasis of size in tis population, but it does substantially reduce the strength of positive directional selection of size that would otherwise occur. While selection currently favours reduced probability of early pregnancy, largely irrespective of body size, it is likely that the occurrence of early pregnancy could result from adaptation to conditions during a recent period during which population density was much lower.

## Introduction

The *paradox of stasis* (Sogard, 1997) and the pattern of *bigger-is-better* with respect to selection on body size have received great attention in the last few decades from both macro- and microevolutionary biologists. Positive directional selection on size appears to be widespread (e.g. Kingsolver & Pfennig, 2004) and genetic variability is generally sufficient for rapid evolution (e.g. Postma, 2014), yet both fossil records (Estes & Arnold, 2007; Hunt, 2007) and extant taxa (Larsson et al., 1998; Knouft & Page, 2003; Ozgul et al., 2009) generally show very little change in body size over time. Both theoretical and methodological explanations have been identified for this mismatch (Bradshaw, 1991; Merila et al., 2001; Hansen & Houle, 2004). One such mechanism, which is expected to be common in nature, is the existence of opposing selective forces (i.e. *trade-offs*), such as those between natural and sexual selection (e.g. Roff, 2000; Roff & Fairbairn, 2007).

Fecundity and sexual selection are major evolutionary forces selecting for larger body size in both females and males (Fairbairn, 1997; Fairbairn et al., 2007). These may be counterbalanced by viability selection against bigger individuals (Blanckenhorn, 2000), leading to a trade-off between sexual (or fecundity) selection and viability selection (Roff, 2000; Roff & Fairbairn, 2007). The benefits and costs associated with larger body size provide the mechanisms underlying such trade-offs and have been extensively discussed (Shine, 1988; Honek, 1993; Anderson, 1995; Sogard, 1997; Blanckenhorn, 2000; Sokolovska et al., 2000). This interplay of costs and benefits is naturally not limited to adult body size. In particular, Blanckenhorn (2000) emphasizes the distinction between the costs of *becoming* as opposed to *being* large, and elaborates on how forces such as predation, parasitism and starvation play different roles in juveniles and adults.

Trade-offs are most readily identified through a deep understanding of a species’ biology. Once a trade-off’s mechanistic basis is hypothesised, multivariate statistical methods can be used to characterise trade-offs in evolutionary quantitative genetic terms. Ultimately, trade-offs result when a character has both positive and negative effects on fitness, acting through different mechanisms. Path analysis, developed by Wright (1921, 1934) to deal “with a system of interrelated variables” (Wright, 1960), is a natural approach for quantifying trade-offs, as it provides the means to disentangle the various mechanisms by which body size can affect fitness. Path analysis underlies the concept of the *extended* selection gradient, formally defined by Morrissey (2014) as the total causal effects of traits on fitness, i.e, taking into account the whole system of causal associations among traits. Extended selection gradients are therefore akin to the concept of *selection for*, defined by Sober (1986) in opposition to that of *selection of* (the total association of traits with fitness). The initial linear formulation of path analysis and extended selection gradients was subsequently extended to non-linear development systems (Morrissey, 2015). Such non-linear systems are typical in studies of selection, which often include variables (e.g. survival probability or offspring number) that are non-linear functions of underlying traits. Recently, Henshaw et al. (2020) provided a means of decomposing the total causal effect of a trait on fitness (i.e. the extended selection gradient) into components associated with multiple non-linear paths.

The Soay sheep (*Ovis aries*) on St Kilda (Clutton-Brock & Pemberton, 2004b) is an example of a system in which the paradox of stasis occurs in relation to body size. Positive directional selection (Milner et al., 2004; Morrissey et al., 2012a; Hunter et al., 2018) and reasonably high heritability (Milner et al., 2000; Wilson et al., 2007; Bérénos et al., 2014) have been reported for body size in this system, alongside stasis (Ozgul et al., 2009). Faster growth may be harmful in juvenile females due to an association between size and early fecundity, combined with a viability cost of reproduction. A non-negligible percentage of ewes get pregnant as lambs and those ewes are more likely to die during their first year of life (Clutton-Brock et al., 2004). We hypothesise that early pregnancy is size-dependent, with larger ewes being more likely to get pregnant and, therefore, to die. A mechanism regulating body size emerges directly from this hypothesis – genetic variation in body mass could be maintained by a trade-off between fitness components in female lambs: being larger is associated with higher rates of survival and total offspring production over a lifetime (Clutton-Brock et al., 1992, 1996; Milner et al., 1999), but being pregnant as a lamb would be associated with lower first-year survival.

In this study, we first model the relationship lamb body mass and early pregnancy, as well as their effects on first-year survival and lifetime offspring production, showing that: (1) early pregnancy is size-dependent, (2) there is a positive additive genetic correlation between lamb body mass and early pregnancy, (3) there is viability selection against early pregnancy, and (4) lifetime rearing success (LReS; lambs reared to independence) in surviving ewes is size-dependent, but very similar in ewes that did and did not become pregnant in their first year. We subsequently draw on these models, in combination with recently developed theory for the analysis and decomposition of selection in non-linear systems (Morrissey, 2015; Henshaw et al., 2020) to implement a formal quantitative genetic analysis of natural selection. We estimate overall selection on body mass to be positive, and we quantify how much stronger it would be in the absence of lamb pregnancy. We also show that about half the selection on lamb body mass occurs later in life, suggesting that this trade-off also regulates body size in adults through across-age correlation in body size.

## Study system

### Data collection

The population of Soay sheep inhabiting the Village Bay on St Kilda in the Outer Hebrides (57°49’N 08°34’W) has been the subject of intensive, individual-based study since 1985 (Clutton-Brock & Pemberton, 2004a). More than 95% of the individuals living in the study area have been marked with plastic ear tags shortly after birth to enable identification throughout their lifetimes, at which time maternal identity and twin status is ascertained from field observations. Regardless of whether a ewe’s lamb is captured (e.g., in cases of neonatal mortality), reproductive status is extensively recorded, allowing assessment of annual and lifetime fitness of ewes. Regular censuses and mortality searches provide death dates for the majority of the sheep. Each year in August a large portion of the study population is captured and phenotyped for multiple traits, including body mass. We focus on phenotypic and life history data on ewe lambs from 1991 to 2015, corresponding to 3457 ewes, as there is no systematic record of pregnancy status from post-mortems of ewes that died during the winter before 1991.

The pedigree of this population has been constructed through a combination of observational field data and molecular markers for maternal links, and using molecular markers only for paternal links (Johnston et al., 2013; Bérénos et al., 2014). 315 polymorphic and unlinked SNP markers were used in molecular parentage assignments (for 4371 individuals) with 100% confidence in the R package MasterBayes (Hadfield et al., 2006). Polymorphic microsatellite markers were also used when SNP genotypes were not available for either lamb or candidate fathers. In those cases, for a total of 222 lambs, 14-18 polymorphic microsatellite markers were used in assignments with confidence >95% in MasterBayes (Morrissey et al., 2012b). The resulting pedigree has a maximum depth of 10 generations and consists of 6740 individuals, of which 6336 are nonfounders (i.e. have one or two known parents).

### Summary statistics of mass and early pregnancy

Early pregnancy, here defined as a pregnancy occurring during the first year of life, occurs at a rate of 37 per year in Soay sheep (914 records documented from 1991 to 2015). The frequency of lamb pregnancy varies greatly among years, but in the raw data no apparent temporal trend is found (Fig. 1a). A major driver of these pregnancies seems to be body size, as the probability of pregnancy increases significantly with body mass (Fig. 1b). A lamb weighing around 10 kg has a very low chance of becoming pregnant, whereas a lamb weighing around 15 kg is more likely to become pregnant than not (Fig. 1c). It is well known that body mass in Soay sheep depends on population size, on average decreasing in years with higher density (Clutton-Brock & Pemberton, 2004b; Ozgul et al., 2009). Conditional on population density, there is a temporal trend of decreasing body mass in Soay sheep, including in lambs (Ozgul et al., 2009), and also in ewe lambs (Fig. 1d, upper panel). This trend is apparently concomitant with a decrease in the rate of early pregnancy over the years, when density is again accounted for (Fig. 1d, lower panel). Substantial costs and few benefits would explain this trend in early pregnancy, as, for example, ewe lamb descendants having significant lower survival when compared to offspring born to older ewes (Fig. 1e).

**Figure 1:**
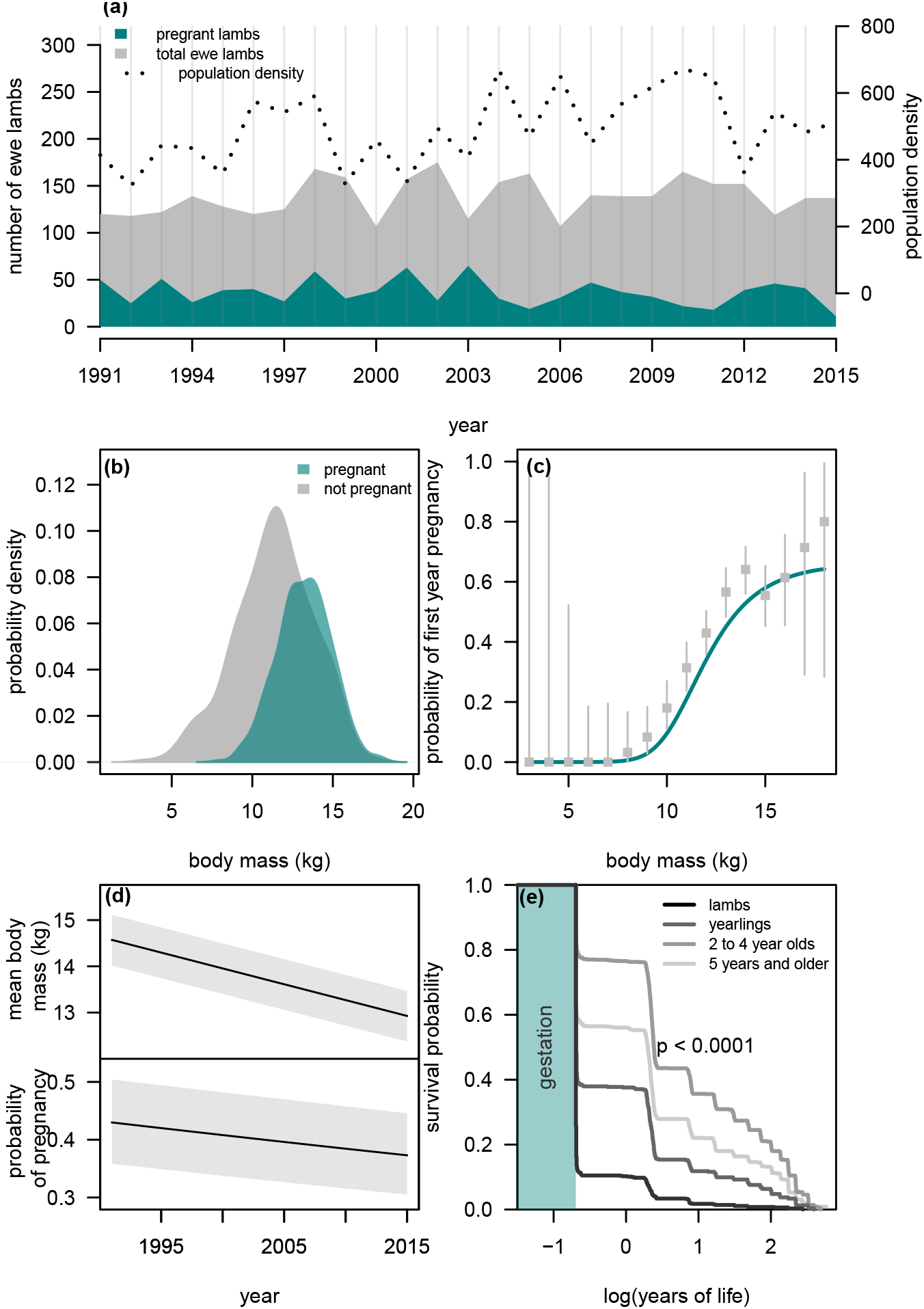
(a) Number of Soay sheep ewes, ewe lambs and population size from 1991 to 2015 in the Village Bay study area; (b) empirical distribution of body mass in pregnant and non-pregnant ewe lambs; (c) size-dependent probability of early pregnancy. The line in teal corresponds to the prediction of a binomial regression with logit link function (see Appendix B for full specification and parameter estimates), whereas the grey bars correspond to 95% confidence intervals on the binomial probability obtained from the raw data; (d) lamb body mass (upper panel) and probability of early pregnancy (lower panel) across time. The black lines correspond to predictions of a linear and a binomial model (logit link function), respectively, correcting for within-year birth and measurement dates, maternal age and population density (linear and quadratic terms for the latter two covariates). Random effects for cohort and maternal identity were also considered. The grey areas illustrate the variation in lamb body mass and probability of early pregnancy attributed to varying population density in 1 standard deviation from the mean; (e) non-parametric survival curves of Soay sheep according to mother’s age at conception. The *p* value corresponds to a Peto & Peto test for differences among Kaplan-Meier curves.

## Models of size, early pregnancy, survival and fitness

In this section, we investigate associations among body size, early pregnancy, survival, and parental success. The purpose of these models is to dissect all of the individual associations among key variables that each contribute to some aspect of overall selection on size and early pregnancy. Ultimately, these models will parameterise the functions relating variables in a path analysis, allowing to disentangling selection *of* and *for* body mass and early pregnancy. All models were fit with MCMCglmm (Hadfield, 2010), using inverse gamma distributions for residual variances (of non-binary traits), and inverse Wishart distributions for the (co)variance components associated with random effects as prior distributions. Also, parameter expansion was implemented to increase convergence efficiency (Gelman, 2006; Hadfield, 2010).

Covariates are included in the models such that fitness and selection are obtained for average conditions of such variables (e.g. maternal age at parity). For clarity, simplified versions of model descriptions, excluding these variables, are presented in the main text, but complete information is provided in Appendix B. Two exceptions are made: twin status (singleton or twin) and population density. In the former case, the binary nature of the variable demands its relationship to the variables included in the path analysis to be explicitly modelled. As for the latter, given the dynamics of the Soay sheep population in St Kilda, understanding how fitness and selection vary with population density is key to understanding selection of body size and pregnancy. All interaction terms are omitted in equations shown in the main text, but are reported in Appendix B.

We define parental success as the number of offspring successfully reared to November 1^st^ and distinguish between those offspring successfully reared as a result of early pregnancy (annual rearing success, AReS) and subsequently (lifetime rearing success, LReS). While these measures of reproductive fitness differ from the traditional definitions of breeding success (offspring produced over a lifetime) and reproductive success (offspring successfully raised up until the age of 1; Clutton-Brock 1988), they delineate fitness associated with a ewe’s ability to gestate and provision a lamb from subsequent components of lamb survival which may be less directly dependent on the mother.

### Lamb body size

A linear regression of the form

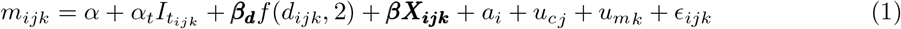

was adopted to model August body mass of lamb *i*, born in year *j* to mother *k*. *α* and *α_t_* are the intercept and the twinning-specific contrast to the intercept, ***β_d_*** are the linear (slope) and quadratic terms associated to a second order polynomial function on population density, *d*, and ***β*** is a vector with the coefficients for the remaining fixed effects, ***X*** (see Appendix B). Random intercepts to estimate among-mother (*u_mk_*) and among-cohort (*u_cj_*) variation were also included. Finally, *α_i_* is the breeding value of lamb *i* and *ϵ* are the residuals of the model. Parameter estimates are provided in Appendix B, Table B1.

### Early pregnancy

The probablibility of early pregnancy was estimated using a binomial regression with a logistic link function,

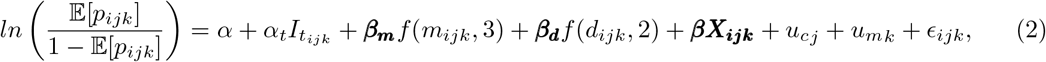

where ***β_m_*** corresponds to a vector with coeffcients for the linear, quadratic, and cubic terms for August body mass, *m*. The remaining quantities have analogous meanings as in Equation (1). More generally, although corresponding to different quantities, we use the same Greek letters for equivalent coefficients in different models throughout the manuscript. Finally, as the overdispersion variance is unobservable for binary data, the variance in *ϵ* was set to one. Parameter estimates are provided in Appendix B, Table B2.

### Bivariate association of pregnancy and size

An important step to disentangling the selection *of* and *for* body mass and early pregnancy is to quantify the genetic association between these variables, a signature of the causal relationships between them (Rausher, 1992; Morrissey et al., 2010). To investigate whether such an association exists, in the form of additive genetic covariance, we used an animal model corresponding to a multi-response generalised linear mixed model (Hadfield, 2010) for body mass, *m*, and the probability of lamb pregnancy, *p*. Although the probability of lamb pregnancy is well described by a binomial distribution, a latent scale variable, *p_l_*, can be defined such that 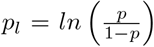. In that case, both *m* and *p_l_* are assumed to be drawn from a multivariate normal distribution, with a mean vector that includes both mean mass, *μ_m_*, and mean pregnancy probability in the logit scale, *μ_p_l__*, and a variance-covariance matrix ***P_l_***,

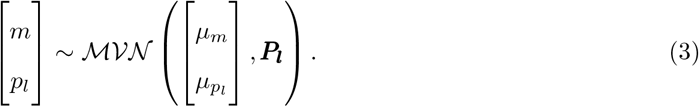

Both *μ_m_* and *μ_p_l__* depend on twin status (singleton or twin), population density, and maternal age at parity, whereas the former also depends on birth and measurement dates. The corresponding parameters and their estimates are listed in Appendix B. Note that the subscript *l* denotes *latent scale*, which here applies to early pregnancy only, since August lamb body mass was modelled on its natural scale. The genetic and environmental contributions to ***P_l_*** were partitioned by including random effects on breeding values (animal model, Henderson 1975). In fact, a ***G_l_*** matrix and an ***E_l_*** matrix can be defined such that

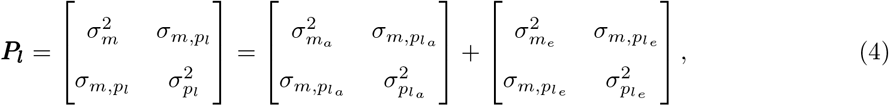

where subscripts *a* and *e* denote the additive genetic and the environmental contribution to the overall (co)variances, respectively. For more information on (co)variance partition, see Falconer (1981). To simplify the notation, we do not use any extra subscript to denote phenotypic (co)variances (see matrix ***P_l_*** in Equation 4). Besides residual variances and covariances, the matrix ***E_l_*** also includes variances and covariances associated with cohort and maternal random effects. Since the overdispersion variance of a generalised linear mixed model (GLMM) is unobservable for binary data, the residual variance for *p_l_* was set to one.

Taking into account the additive genetic and the phenotypic variances in lamb body mass, 0.78 (95% CrI 0.20; 1.27) and 4.29 (95% CrI 3.84; 5.23), respectively, the heritability of this trait is estimated to be 0.14 (95% CrI 0.04; 0.29), very similar to the estimate reported by Bérénos et al. (2014) for both sexes (0.12, SE 0.036). Up until very recently, for binomial variables, heritabilities were typically obtained by approximation, as described in Nakagawa & Schielzeth (2010). Following this approach, we obtain an estimate of 0.15 95% CrI 0.10; 0.70) for the heritability of early pregnancy. There is a strong positive phenotypic correlation between lamb body mass and early pregnancy (0.51, 95% CrI 0.40; 0.66), the genetic contribution to that correlation being very important (0.64, 95% CrI 0.24; 0.90).

In 2016, Villemereuil et al. derived exact expressions to convert the latent scale parameters estimated in GLMMs into scales on which traits are expressed and selected. We are particularly interested in obtaining parameters in the probability scale (probability of early pregnancy), rather than in the logarithm of the odds of being pregnant as a lamb. Villemereuil et al. (2016) refer both to *expected* and *data* scales, the difference between them being the random noise that distinguishes a model from the data itself. Here, we adopt the expected scale. Following their work, the expected values for body mass and the probability of early pregnancy is given by

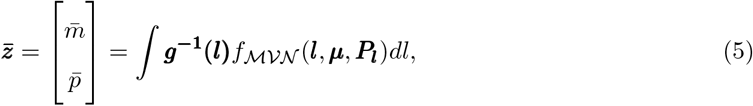

where ***g***^−1^ corresponds to the inverse link functions used to model body mass and the probability of early pregnancy. As the adopted link functions for these variables were the identity and the logit functions, respectively, their inverses are the identity and the logistic functions, the latter corresponding to 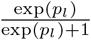. ***l*** are the latent values for lamb body mass and early pregnancy. 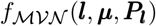 is the probability density of a multivariate normal distribution with vector mean ***μ*** and variance ***P_l_***. Likewise, the phenotypic variance-covariance matrix on the expected scale is given by

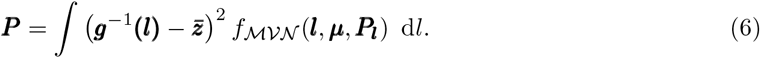

The ***G*** matrix on the expected scale,

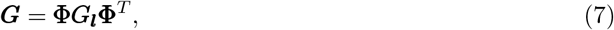

can also be derived (Villemereuil et al., 2016), where **Φ** is the average derivative of the expected values with respect to the latent values,

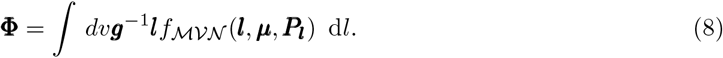

Details on how derivatives and integrals were solved are provided in Appendix C. The heritability on the expected scale is obtained directly through its definition, as the proportion of the additive genetic variance relative to the phenotypic variance, using the derived values. The results obtained using the above expressions can also be found in Table B3. The heritability of early pregnancy on the expected scale (probability) is 0.40 (95% CrI 0.11; 0.63), higher than the approximation shown above (but note that this approximation is derived for the data scale), and corroborating the genetic basis of this trait. Using the derived values on the expected scale, we also obtained the conditional genetic variance of lamb body mass. This measure allows us to evaluate the proportion of the additive genetic variance in lamb body mass that is independent of early pregnancy. Following Hansen et al. (2003), this quantity, 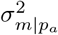, is given by

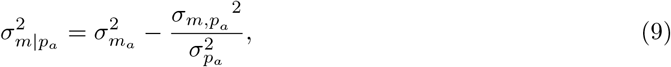

which in this particular case equals 0.40 (95% CrI 0.06; 0.85). The analogous metric for early pregnancy corresponds to 0.03 (95% CrI 0.00; 0.05). Since 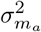 and 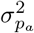 were estimated to be, respectively, 0.78 (95% CrI 0.20; 1.27) and 0.04 (95% CrI 0.00; 0.08), the additive genetic variance in lamb body mass and early pregnancy that are independent from one another corresponds to 59% (95% CrI 29%; 100%). Substantial proportions of the genetic variances in lamb body mass and early pregnancy are not independent, and therefore *selection for* each one of these traits necessarily results in *selection of* the other.

### First-year survival

We investigated whether early pregnancy has fitness costs by evaluating first-year survival in ewe lambs as a function of pregnancy and body mass. First-year survival was defined using the first day of May as the cut-off date to make sure that dying during labour was considered in the first annual cycle of the ewes (89% of births are before 1 May). Ewes with known first-year survival status and that survived until the rut were included in the analysis (*n*_2_ = 947). A binomial regression with logit link function of the form

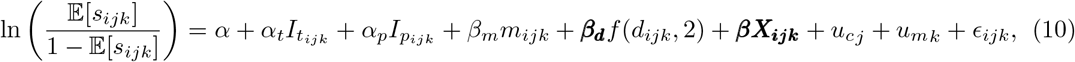

was adopted to model the probability of lamb *i*, born in year *j* to mother *k*, survives its first year of life. *α* and *α_p_* are the intercept and the pregnancy-specific contrast to the intercept, *β_m_* is the slope for lamb body mass, and ***β*** is a vector with the coefficients for the remaining fixed effects. Although not pictured in Equation (10), a pregnancy-specific contrast to the slope for mass is also estimated (Appendix B). As first-year survival, and survival in general, is highly dependent on cohort effects due to variability in winter conditions and food availability (Clutton-Brock et al., 2004), we included cohorts as random effects (*u_cj_*), as well as maternal environmental effects (*u_mk_*). Variance in residuals (*ϵ_ijk_*) was set to one, as overdispersion is unobservable in binomial mixed models.

We first establish that first-year survival is size-dependent by adopting a similar model to the one in Equation (10), but excluding the pregnancy-specific contrasts - bigger ewes have higher chance of survival (*p* < 0.001, Fig. 2a, Tab. B4a in App. B). Adopting the full model in Equation (10) shows that for a given body mass, pregnant lambs are significantly less likely to survive their first annual cycle when compared to the ewes that do not get pregnant (*p* < 0.001, Fig. 2c, Tab. B4b in App. B). Lamb pregnancy has a substantial survival cost - the probability of survival for ewe lambs of average body mass that did not get pregnant is 50% (95% CrI 37%; 65%), an estimate that drops to 26% (95% CrI 14%; 39%) in pregnant lambs. Pregnant lambs of all sizes are more likely to die than non-pregnant ewe lambs (Fig. 2c). Although merely a numerical result, it is interesting to notice that mean body mass is lower among *all* lambs that got pregnant (14.00, 95% CrI 13.65; 14.48) than it is in the sub-group of lambs that got pregnant and survived at least a year (14.59, 95% CrI 14.11; 15.03), suggesting that if a ewe is to be large enough to get pregnant, then it is better to be as large as possible. Investigating the simultaneous effect of lamb body mass, early pregnancy and population density on first-year survival exposes an important interaction effect between the former traits and the latter. Although large population size is, in general, associated with lower survival, it is evident that pregnant ewe lambs are more susceptible to variation in population density than lambs that do not get pregnant. First-year survival in pregnant ewes is very low (below 10%) across most possible values of population density, unless body mass is relatively very high, whereas such low yearly survival rates are much less likely in non-pregnant ewes (App. A, Fig. A1a,b).

**Figure 2:**
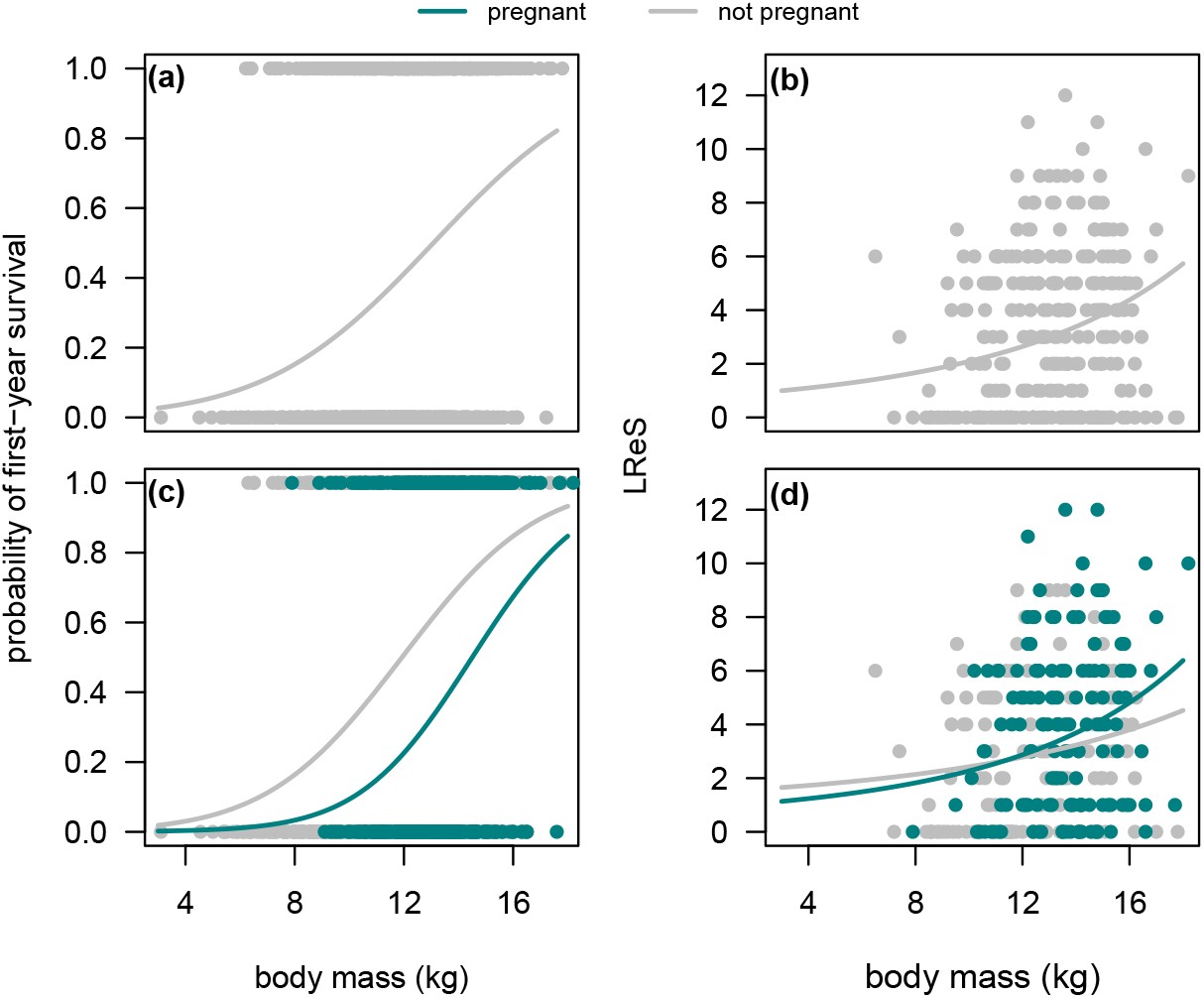
Probability of first-year survival (a, c) and lifetime rearing success, LReS, (b, d) in Soay sheep ewes as a function of lamb body mass (a, b), and both lamb body mass and early pregnancy status (c, d). Dots correspond to observed data and curves to model predictions. Binomial regressions for first-year survival with logit link function were fit to the data. LReS was estimated in ewes that survived their first year of life and Poisson regressions with exponential link function were fit to the data. In all models, population density and maternal age were included as covariates (linear and quadratic terms were estimated for both), as well as birth and measurement dates (see Tab. B4 and B6 in App. B). Variance among cohorts and maternal identities were also estimated.

### First-year rearing success (AReS)

Having established that pregnant ewe lambs are more likely to die when compared to same-sized non-pregnant ewe lambs, it is important to understand the fate of those pregnancies. From the 914 documented early pregnancies, only 97 resulted in offspring surviving until the winter (AReS), and only 36, less than 5%, resulted in recruitments to the population, i.e. offspring that survived until their first April 1^st^. We estimated the AReS of ewe lamb *i* fitting a binomial regression with a logit link function of the form

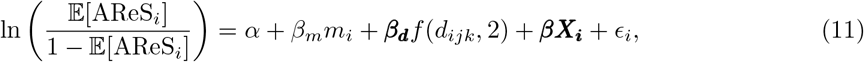

where *α* is the model intercept, *β_m_* is a slope for lamb body mass, and ***β*** is a vector with the coefficients associated to the covariates in ***X***, which includes capture and birth dates, as well a linear and a quadratic terms for population density. The variance in the model residuals, *ϵ_i_*, was set to one. Due to sample size limitations, we did not consider any random effects.

Only 2.2% (95% CrI 0.57%; 6.17%) of the offspring born to ewes getting pregnant as average-sized lambs in average-density years survive until the winter (App. B, Tab. B5). It is interesting to note that the slope for body mass is positive and significantly different from zero, showing that larger ewe lambs are more likely to have surviving offspring, and suggesting that, as for first-year survival, if a ewe lamb is to get pregnant, than the larger the better. Overall, AReS not only increases with ewe body mass, but also significantly decreases as population size increases (App. A, Fig. A1c).

### Subsequent lifetime rearing success (LReS)

We also investigated the effect of lamb body mass and early pregnancy on LReS. As the LReS of ewes dying during their first year of life is zero, analyses presented here only include ewes that survived their first winter. The LReS of ewe *i*, born in year *j* to mother *k* was modelled with a Poisson mixed model with a log link function of the form

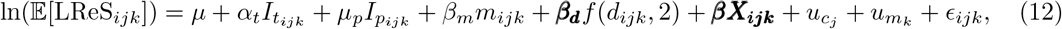

where *μ* is the model intercept and *μ_t_* and *μ_p_* are the twin status and early pregnancy specific contrasts to the intercept, respectively. *β_m_* is the slope for lamb August mass, ***β_d_*** the linear (slope) and quadratic terms for density, and ***β*** a vector with the slopes of the remaining fixed effects, ***X***. Although not shown, a pregnancy-specific contrast to the slope for mass is also estimated (Appendix B). ***u_c_*** and ***u_m_*** are random effects, assumed to be drawn from independent normal distributions, for cohort and maternal identity. Finally, *ϵ_ijk_* is the residual of individual *i*. To study LReS, the data were further constrained to only include ewes from cohorts that were completely phenotyped for this trait, which was achieved by excluding all ewes born after 2006 (*n*_3_ = 550). This procedure resulted in including very few LReS records associated with ewes that were still alive by the cut-off year (2015). While these surviving ewes’ lifetime fitness is underestimated, they have reared most the offspring they will rear in their lives. Excluding these records would result in stronger bias because ewes that lived longer had potentially higher rearing success.

We applied the model in Equation (12) without the coefficients associated with early pregnancy and established that LReS is dependent on lamb size (Fig. 2b, Tab. B6a in App. B). Rearing success is an increasing function of lamb August body mass, with bigger animals being more likely to rear more offspring. Results from the full model suggest that rearing success is not determined by early pregnancy status (Fig. 2d, Tab. B6b), although it is unclear whether under particular conditions of population density this trait could have an effect on LReS that is independent from its correlation with lamb body mass. Neither the contrast to the intercept (0.37; 95% CrI −0.06; 0.82) nor the interaction terms with body mass (0.02; 95% CrI −0.05; 0.07) and population density (3.4 × 10^−3^; 95% CrI −9 × 10^−3^; 7.5 × 10^−3^) differ from zero, however most of the posterior density of the pregnancy-related coefficients supports positive values. Such numeric differences result in distinct expected numbers of offspring successfully reared in ewes that were and were not pregnant as lambs. In averaged-sized individuals living under average environmental conditions, 2.48 (95% CrI 1.57; 4.00) offspring are expected to be reared by ewes that survived their first year of life, which is broken down to 3.36 (95% CrI 1.87; 5.54) and 2.30 (95% CrI 1.40; 3.46) in ewes that got and did not get pregnant as lambs, respectively (Fig. 2b). As no strong statistical support is found for these differences, nor is any biological justification known for such an effect of early pregnancy, LReS will be assumed to be independent from this trait in further analyses. The simultaneous effect of body mass and population density on LReS, independent of early pregnancy, is presented in Appendix A (Fig. A1d), showing not only that LReS is highest in ewes born at low population densities, but also that the effect of lamb body mass on LReS is stronger in ewes that were also born in years of low population density.

### Twinning

The parameterisation of the path model built in this study requires knowledge of the effects of twin status on size, pregnancy, survival, and rearing success, which are estimated in Equations (1), (2), (10), and (12). Furthermore, given the binary nature of the variable, a model of the probability of being a twin is required so that the quantities such as mean phenotype or selection gradients can be average over this probability. For this purpose, a binomial mixed model was fitted according to

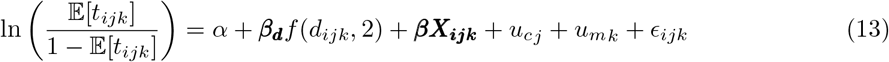

where 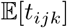 are individual probabilities of twinning, and other terms are defined analogously to other models. Detailed information on the estimated quantities is provided in Appendix B, Table B7.

## Selection analysis

The analyses shown so far establish that (1) early pregnancy is size-dependent, (2) there is a fairly strong positive additive genetic correlation between lamb body mass and early pregnancy, (3) there is selection against early pregnancy through viability selection, (4) the ultimate fitness benefit of early pregnancy is very small, and (5) subsequent LReS in surviving ewes is size-dependent, but very similar in ewes that got and did not get pregnant. In this section, we use the information presented so far in a formal selection analysis to investigate *selection for* and *selection of* lamb size in Soay sheep ewes. In particular, we use a path analysis involving all pertinent traits (Fig. 3), allowing the estimation of an *extended selection gradient* (Morrissey, 2014) for lamb body mass. Using the path rules for developmental systems set by Wright (1934) and expanded to non-linear variables by Morrissey (2015) one can use the twin model approach developed by Henshaw et al. (2020) to decompose the components of the effect of lamb body mass on fitness and that arise via early pregnancy and otherwise. The path diagram shown in Figure 3 includes twining, *t*, mass, *m*, pregnancy, *p*, survival, *s*, first-year rearing success, AReS, subsequent rearing success, LReS, and fitness, *W*, and can be represented by a vector-valued function of the following form

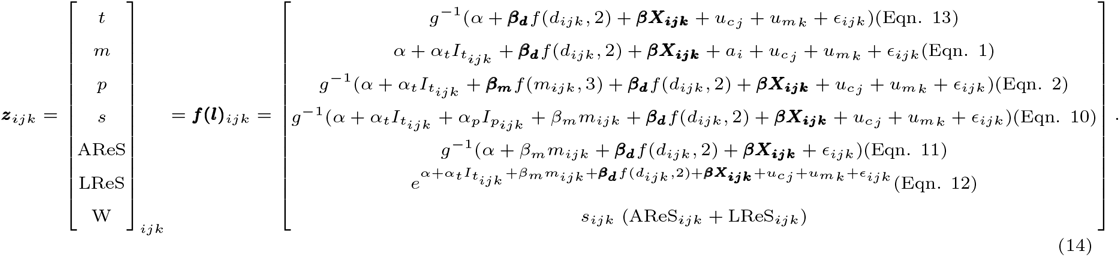

**Figure 3:**
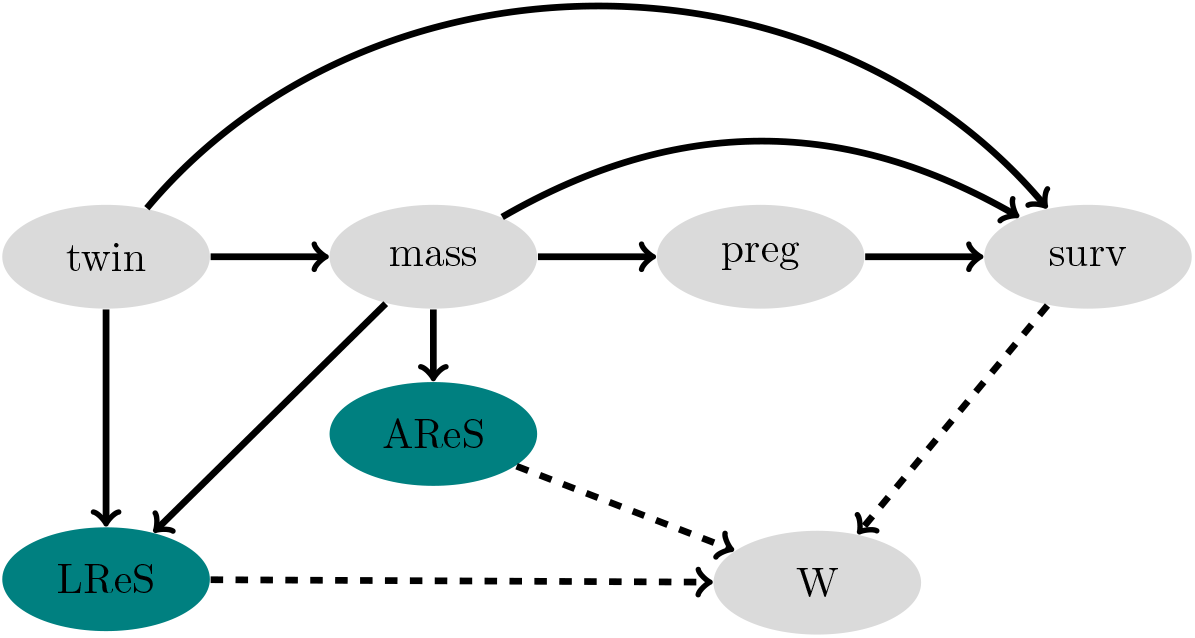
Path diagram representing the phenotypic landscape defined in Equation (14). The probability of twinning, *twin*, has an effect on all the other traits, lamb body *mass* affects the probabilities of being pregnant as a lamb, *preg*, surviving the first annual cycle, *surv*, and also first year and subsequent lifetime rearing success, *AReS* and *LReS*; and *preg* affects *surv*. Fitness, *W*, is defined as the sum of AReS and LReS on surviving ewes weighted by the probability of first-year survival.

Equation umbers given in parentheses reference the statistical models used to parameterise each component. As before, although corresponding to different quantities, we use the same Greek letters for equivalent coefficients in all models: *α* represents the intercepts, *α_t_* and *α_p_* the twinning and pregnancy-specific contrasts to the intercept, respectively, *β_m_* and ***β_m_*** the slope and the linear (slope), quadratic and cubic terms for mass, ***β_d_*** the linear and quadratic terms for population density and, finally, ***β*** is a vector containing the effects associated with the remaining covariates. ***β*** also includes interaction terms for body mass and population density, interaction terms between early pregnancy status and population density, and between these two and body mass, in all models where the respective predictors are included. *u_cj_* and *u_mk_* are random effects associated with cohort *j* and mother *k*, and *ϵ_ijk_* is the residual of individual *i*. The model for mass also includes breeding values, *a*, allowing us to segregate the additive genetic variance from other sources of variation. Twinning, pregnancy, survival, and AReS were assumed to follow binomial distributions, mass to follow a Gaussian distribution, and LReS to follow a Poisson distribution with additive overdispersion. Fitness was defined as the sum of AReS and LReS of ewes surviving their first annual cycle, weighted by the probability of first-year survival. Note that although population density is not included in the path diagram of Figure 3, all the results presented in this section were obtained by marginalising over the observed population density in all cohorts. Equation (14) was evaluated for the values of population density of each cohort and the mean value taken as the best estimate of 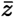.

### Selection on mass and the probability of pregnancy

Mean phenotype, 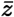, and the phenotypic variance-covariance matrix, **P**, on the expected scale were obtained using Equations (5) and (2). We estimate 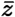 as

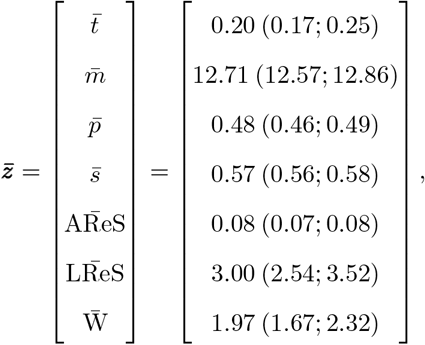

and **P** as shown in Table 2. As variance in breeding values was only estimated for lamb body mass, which was modelled on its natural scale, the additive genetic variance in mass on the expected scale is the one estimated in the model, 0.67 (95% CrI 0.07; 1.14). The fact that the correlations between lamb body mass and first-year survival (0.42, 95% CrI 0.33; 0.51) and between early pregnancy and first-year survival (0.11, 95% CrI 0.03; 0.19) are both positive suggests that the equilibrium between lamb body mass affecting first-year survival positively (direct effect) and negatively (through its effect on early pregnancy) is leaning towards the former (Tab. 2). To investigate the consequences of this trade-off in the selection of lamb body mass, we calculated its *extended* directional selection gradient. For trait *z*, the extended directional selection gradient is defined as the average derivative of expected fitness with respect to latent value (Morrissey, 2015; Villemereuil et al., 2016),

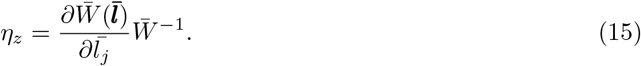

**Table 1:** Coefficients on the latent and expected scales of the multi-response animal model of lamb body mass and early pregnancy in Soay sheep. Lamb mass was modelled on its natural scale, and therefore no distinction between latent and expected scales is necessary. Under the binomial response model for pregnancy, variance components are not meaningful on the expected scale, though means and correlations are. Fixed effects from this model are reported in appendix B.

|  | latent scale |  | conversions to the expected scale |  |
| --- | --- | --- | --- | --- |
|  | posterior mode | 95% CrI | posterior mode | 95% CrI |
| $\mu_m$ | 13.82 | (12.22; 14.26) | | |
| $\mu_p$ | 0.00 | (-0.85; 0.64) | 0.50 | (0.39; 0.59) |
| $\sigma_{m_a}^2$ | 0.78 | (0.20; 1.27) | | |
| $\sigma_{p_a}^2$ | 0.69 | (0.00; 5.86) | | |
| $\sigma_{mp_a}$ | 0.31 | (0.00; 1.49) | | |
| $\rho_{mp_a}$ | 0.64 | (0.24; 0.90) | 0.64 | (0.24; 0.90) |
| $\sigma_m^2$ | 4.29 | (3.84; 5.23) | | |
| $\sigma_p^2$ | 3.20 | (1.99; 10.12) | | |
| $\sigma_{mp}$ | 3.24 | (1.49; 3.50) | | |
| $\rho_{mp}$ | 0.51 | (0.40; 0.66) | 0.49 | (0.38; 0.64) |

**Table 2:** Phenotypic variance-covariance matrix for the path diagram in Fig. 4, including the probability of twinning (*t*), August body mass (*m*), the probability of early pregnancy (*p*), the probability of survival (*s*), first year rearing success (*ares*), subsequent lifetime rearing success (*lres*), and fitness, defined as lifetime rearing success (AReS+LReS) in surviving ewes weighed by the probability of survival. The diagonal corresponds to variances, the upper off-diagonal to covariances and the lower off-diagonal to correlations among traits. Values shown correspond to the expected scale.

|  | t | m | p | s | ares | lres | W |
| --- | --- | --- | --- | --- | --- | --- | --- |
| t | 0.09 (0.06; 0.11) | -0.25 (-0.34; -0.16) | -0.01 (-0.02; -0.01) | -0.02 (-0.02; -0.01) | 0.00 (0.00; 0.00) | -0.05 (-0.13; 0.01) | -0.09 (-0.15; -0.04) |
| m | -0.43 (-0.49; -0.34) | 3.92 (3.54; 4.55) | 0.31 (0.27; 0.36) | 0.25 (0.20; 0.29) | 0.04 (0.03; 0.05) | 1.26 (0.86; 2.02) | 1.45 (1.17; 2.18) |
| p | -0.14 (-0.22; -0.08) | 0.57 (0.48; 0.64) | 0.08 (0.07; 0.09) | 0.01 (0.00; 0.02) | 0.00 (0.00; 0.00) | 0.10 (0.04; 0.18) | 0.09 (0.04; 0.15) |
| s | -0.21 (-0.27; -0.13) | 0.42 (0.33; 0.51) | 0.11 (0.03; 0.19) | 0.09 (0.07; 0.11) | 0.00 (0.00; 0.00) | 0.08 (0.03; 0.16) | 0.31 (0.23; 0.40) |
| ares | -0.12 (-0.17; -0.06) | 0.31 (0.25; 0.36) | 0.15 (0.09; 0.21) | 0.14 (0.08; 0.19) | 0.00 (0.00; 0.01) | 0.01 (0.00; 0.04) | 0.02 (0.01; 0.04) |
| lres | -0.05 (-0.11; 0.00) | 0.19 (0.11; 0.26) | 0.10 (0.03; 0.16) | 0.08 (0.02; 0.13) | 0.05 (-0.01; 0.13) | 11.39 (4.93; 36.23) | 8.51 (3.74; 25.96) |
| W | -0.09 (-0.15; -0.05) | 0.27 (0.18; 0.35) | 0.10 (0.03; 0.16) | 0.34 (0.24; 0.43) | 0.11 (0.03; 0.18) | 0.87 (0.79; 0.93) | 7.55 (3.88; 23.60) |

This selection gradient thus represents the ultimate effect of small changes in phenotype (possibly, latent values for phenotypes that are modelled probabilistically, as is pregnancy) on fitness, scaled to relative fitness, averaged over the range of conditions experienced by individuals in the population. The algorithm for implementing the calculation described by equation 15 is detailed in appendix C.

The extended selection gradient for mass is positive, *η_m_* = 0.26 (95% CrI 0.25; 0.28), whereas for lamb pregnancy the extended selection gradient is negative, *η_p_* = −0.56 (95% CrI −0.64; −0.47). Importantly, the conflict between viability and fecundity is density-dependent, and is almost nonexistent when population density is very low. In such circumstances, virtually all ewe lambs survive, and therefore early pregnancy does not have a significant cost. As a consequence, *η_p_* is higher (i.e., less strongly favouring low values) when population density is at its lowest and decreases with population density (Fig. 4, Tab. 3). On the other hand, at high densities, when the probability of first-year survival is lowest, selection favouring larger body size is stronger.

**Figure 4:**
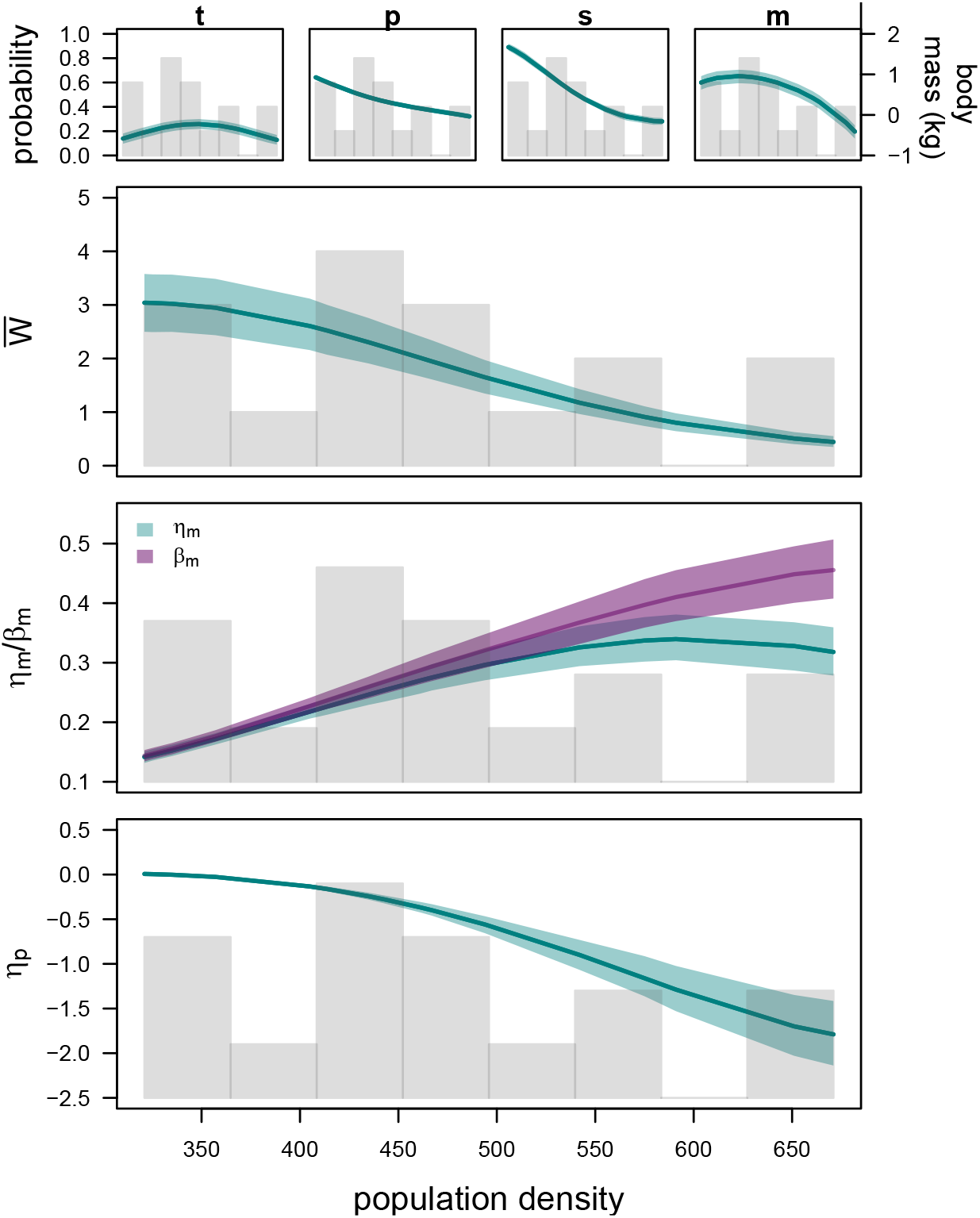
The effect of population density on traits depicted in the path diagram in Figure 3, and consequently on selection of lamb body mass and early pregnancy. The upper panel shows variation in mean twinning (t), early pregnancy (p), first-year survival (s), and mean-centred lamb body mass (m). The three lower panels include functions of mean fitness, selection of lamb body mass (*η_m_* and *β_m_*), and selection of early pregnancy (*η_p_*) on population density. All plots include an histogram showing the empirical distribution of population density. Areas in teal and purple correspond to 95% highest posterior density credible intervals.

**Table 3:** Selection gradients of lamb body mass and early pregnancy for three different values of observed population density - minimum, mean and maximum (321, 452, and 671, respectively). Raw (*η*) and standardised selection gradientes, either by the observed standard deviation (*η_sd_*) or the observed mean (*η_μ_*), are shown. For lamb body mass, selection gradients considering first-year survival as a measure of fitness are also included (*η_s_*).

|  |  | d=321 | d=452 | d=671 |
| --- | --- | --- | --- | --- |
| mass | $\eta$ | 0.15 (0.14; 0.16) | 0.28 (0.27; 0.30) | 0.33 (0.30; 0.36) |
| | $\eta_{sd}$ | 0.35 (0.33; 0.36) | 0.66 (0.62; 0.70) | 0.76 (0.69; 0.84) |
| | $\eta_\mu$ | 1.86 (1.77; 1.96) | 3.53 (3.31; 3.76) | 4.11 (3.71; 4.50) |
| | $\eta_s$ | 0.04 (0.04; 0.05) | 0.15 (0.13; 0.17) | 0.13 (0.11; 0.16) |
| | $\eta_{s_{sd}}$ | 0.10 (0.09; 0.11) | 0.34 (0.30; 0.38) | 0.31 (0.25; 0.37) |
| | $\eta_{s_\mu}$ | 0.53 (0.46; 0.59) | 1.83 (1.62; 2.06) | 1.66 (1.37; 1.97) |
| pregnancy | $\eta$ | 0.01 (0.00; 0.02) | -0.40 (-0.45; -0.34) | -1.79 (-2.08; -1.48) |
| | $\eta_{sd}$ | 0.00 (0.00; 0.01) | -0.20 (-0.22; -0.17) | -0.89 (-1.04; -0.74) |

To investigate the effect of pregnancy on the total effect of mass on fitness, we calculated a direct selection gradient, *β_m_*, as opposed to the *extended* directional selection gradient, *η_m_*. The direct selection gradient describes that part of the effect of a focal trait (mass) on fitness that occurs independently of other focal traits (in this case, pregnancy). We calculated the direct selection gradient using the twin model approach described in, which amounts to re-calculating the selection gradient for mass, but using a modified path diagram (as Figure **??**), in which causal links from the focal trait to other traits (in Figure **??**, this is the effect of mass on pregnancy) are omitted. *β_m_* is thus blind to the positive association that exists between lamb body mass and early pregnancy, and therefore to pregnancy-mediated viability selection on body mass. On average direct and total effects of mass on fitness do not differ much, and *β_m_* is only marginally larger (0.29; 95% CrI 0.27; 0.30). The reason for that lies again in the dependency of these quantities on population density. *η_m_* and *β_m_* are very similar when population density is very low, as viability selection tends to zero and, therefore, there is no cost to early pregnancy. Indeed, we show that *η_m_* and *β_m_* do diverge substantially with increasing population density (Fig. 4, Tab. 3), indicating that the mediation of selection for mass by pregnancy is highly environment-dependent.

The path analysis represented by Figure 3 invokes a definition of fitness that includes the entire life cycle. However, the mechanisms through which selection acts, survival and fecundity, occur at different points of the life cycle. Viability selection is particularly strong in the first annual cycle, whereas most offspring are reared at older ages. To evaluate the strength of the two mechanisms is, therefore, also to understand in which period selection is the strongest. We quantify the contribution of selection on lamb body mass that occurs during the first year of life using a path diagram similar to the one in Figure 3, but defining fitness as first-year survival. Measured this way, *η_m_* is 0.12 (95% CrI 0.11; 0.13), and therefore corresponds to about half of the total lifetime selection on body mass. Selection acting on lamb body mass through viability costs during the first year of life is proportionally very low when population density is lowest (around 30%), is highest at intermediate populations densities and decreases when population density is very high. A slightly different pattern is obtained if *β_m_*, instead of *η_m_*, is used to quantify the strength of selection. In this case, the relative importance of viability selection does not decrease when population density is at its highest. These two trajectories, put together, suggest that at low densities, when both fecundity and survival are very high (Fig. 4), fecundity governs selection on lamb body mass, as mothers will successfully rear several offspring. As population size increases and first-year survival decreases (Fig. 4), viability selection becomes proportionally more important as it occurs before any opportunity for reproduction. Under exceptional high population density, the probability of survival is very low. As most ewes die, fecundity selection becomes slightly more important. That is not observed when the correlation between lamb body mass and early pregnancy is dismissed, because the pregnancy-related viability costs are not considered.

### Selection of the probabilistic reaction norm for early pregnancy

Finally, we investigated in more detail the conditions under which early pregnancy could be advantageous, or at least not maladaptive, to Soay sheep ewes. Knowledge of the selection pressures acting on different components of reaction norms of body mass, along with their potential to evolve, could shed light on the persistence of early pregnancy despite its large viability cost. In this context, we assess whether there is genetic variation in, and selection acting on, the slope of the reaction norm for pregnancy across differences in body mass and environment (population size). First, to quantify genetic variance, we fit a random regression animal model of the form

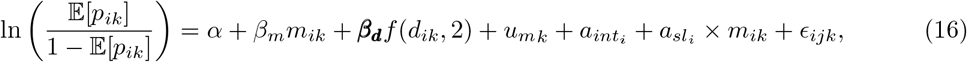

a version of Equation (2) where variance in breeding values is estimated, associated with both the intercept (*a_int_*) and the slope for body mass (*a_sl_*). As for assessing selection, we (1) evaluated the fitness of pregnant and non-pregnant ewes as a function of size and population density, thus obtaining *conditional individual fitness functions*, and we (2) calculated extended selection gradients for the intercept and slope for mass across different contexts of mean phenotype and environment. In (1) we were particularly interested in assessing whether the *conditional individual fitness functions* derived for pregnant and non-pregnant ewes would cross, as that would be a clear indication of variation in selection. In both analyses, we evaluated Equation (14) at fixed values of body mass and population density, setting the probability of pregnancy to 0 or 1 in analysis (1).

Fitting the random regression animal model to estimate additive genetic reaction norms of early pregnancy on body mass was numerically very challenging, both due to the binary nature of the response variable and the lack of repeated measures. Nonetheless, we found strong evidence of additive genetic variance in both the intercepts (2.04, 95% CrI 0.13; 9.19, App. B) and slopes (0.19, 95% CrI 0.05; 0.67) of the random regression being significantly different from zero, confirming a genetic basis of the reaction norms. Additionally, a positive additive genetic covariance between intercepts and slopes also seems to exist (0.61, 95% CrI 0.15; 2.35). On the other hand, we found very little evidence of variation in selection acting on the slope for body mass (Fig. 6). Most conditional individual fitness functions for pregnant and non-pregnant ewes evaluated across a range of population densities never cross, suggesting that the effect of body mass on fitness does not differ qualtatively between pregnant and non-pregnant ewes. Although some of these functions cross, visual inspection suggests that this happens only when functions for pregnant and non-pregnant lambs are very similar (Fig. 6a). Finally, across sensible values of mean phenotype and environmental conditions, estimates of *η* for the slope of the reaction norm of early pregnancy on August mass are very close to zero (Fig. 6b), providing little evidence for variation in selection for plasticity of pregnancy.

**Figure 5:**
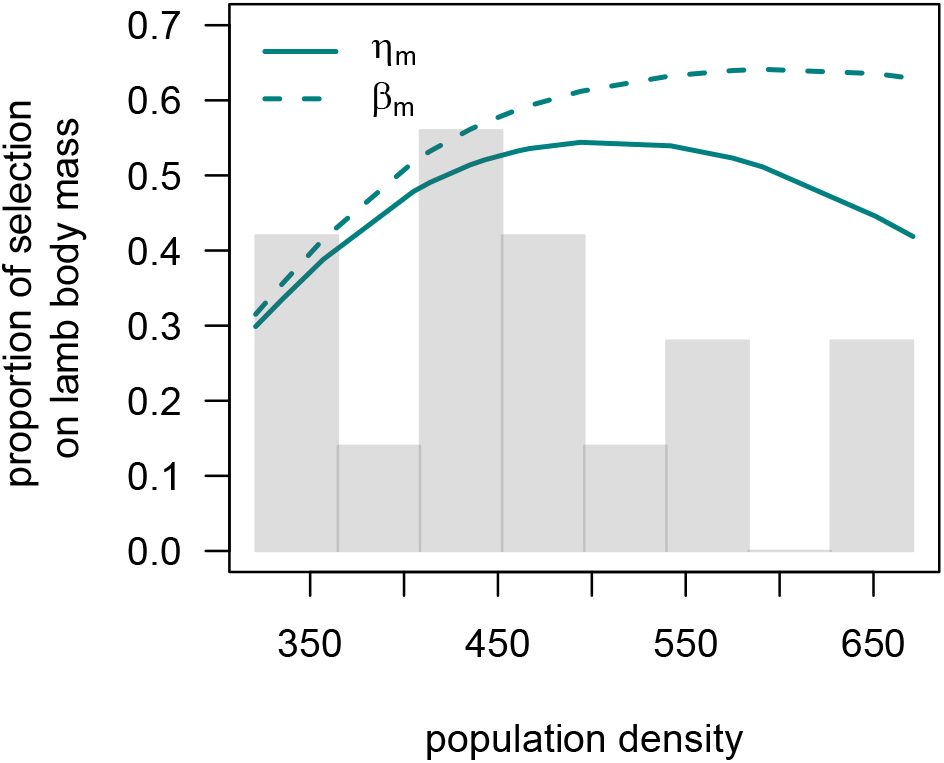
Proportion of natural selection on lamb body mass occurring through viability selection in Soay sheep ewes as a function of population density. This proportion was obtained both using the extended (*η_m_*) and non-extended (*β_m_*) directional selection gradient of lamb body mass. The histogram in grey shows the empirical distribution of population density.

**Figure 6:**
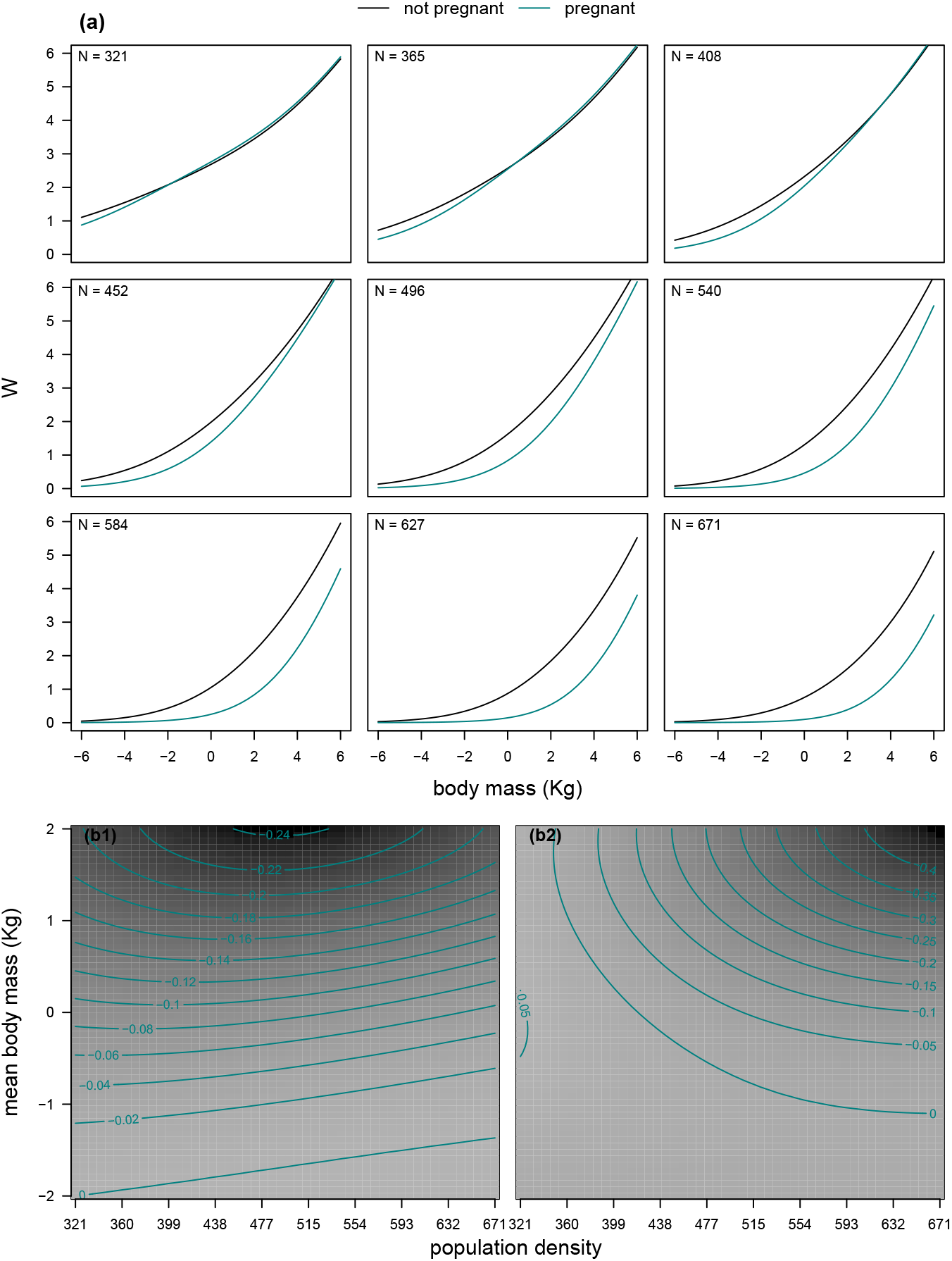
(a) Conditional individual fitness functions of (mean) August body mass for pregnant and non-pregnant Soay sheep ewes and (b) selection gradients for the intercept (b1) and slope of early pregnancy on body mass (b2) as a function of population density and body mass. Body mass is mean-centred in both (a) and (b). Note that if the effect of body mass on the strength of selection varies between pregnant and non-pregnant ewes the conditional individual fitness functions in (a) are expected to cross.

## Discussion

We have characterised a trade-off between viability and fecundity selection of body size in female Soay sheep lambs. Larger ewe lambs are more likely to become pregnant during their first annual cycle, and pregnant lambs are more likely to die. As a consequence, natural selection acts on lamb body mass through two distinct pathways: through the direct effect of lamb body mass on fitness, independent of early pregnancy status, and through its effect on early pregnancy. Selection on lamb body mass would, therefore, be stronger (more positive) if not for the relationship between this trait and early pregnancy. As a result, this correlation is one mechanisms constrains the evolution of body size in Soay sheep.

Recent theory on non-linear development systems allows one to decompose selection across its component causal pathways, essentially by using a system of path analyses that allows for non-linear effects and interactions (Morrissey, 2015; Henshaw et al., 2020). We used this theory to disentangle and quantify the direct effect of lamb body mass on fitness and that arising via its effect on early pregnancy. Alternatives to this approach are the use of traditional path analysis (developed for linear traits, e.g. Scheiner et al., 2000; Morrissey, 2014) or multi-response (linear or generalised linear) models that explicitly estimate (additive genetic) correlations among traits (e.g. Bonnet et al., 2017). Our approach specifically harnesses the phenotypic relationship between lamb body mass and early pregnancy, and builds it into a formal model of selection. Such an approach has the potential to be particularly useful when analytical power is a limiting constraint to infer the genetic architecture among traits. If genetic trade-offs are indeed a major explanation for the widespread mismatch between predicted evolutionary change and observed dynamics of body size (Blanckenhorn, 2000; Hansen & Houle, 2004; Kruuk et al., 2008), why are these methods not more frequently adopted? Information available suggests that the most considerable obstacle in dealing with the selection of correlated traits is not purely methodological. Rather, detecting genetic correlations seems to be a limiting step, with evidence for genetic constraints in the wild being rather scarce (Kruuk et al., 2008). Reasons for this may include insufficient knowledge about the biology and ecology of a species (Pemberton, 2010), but also the presence of confounding effects between genes and environment. Negative genetic covariation between two traits might be obscured at the phenotypic level by environmental covariation among traits (Roff & Fairbairn, 2007; Morrissey et al., 2012c). Put together, evidence so far supports the hypothesis that the paradox of stasis generally has no single and simple solution and that a deep understanding of the species’ biology may be critical to picking apart the causes of associations between traits and fitness.

Assessing the impact of early life events on fitness is often difficult because it requires data collection protocols that are not possible in all study systems. Nonetheless, the characterisation of individuals that do not survive is crucial in estimating evolutionary parameters. As an example, failing to identify and to characterise Soay sheep ewes that die due to pregnancy in their first year of life would have led to overestimating the strength of selection acting on August body mass. Essentially, this is because survival data would be due to data *missing not at random*: an invisible fraction of the population would have not been taken into account. Given that viability selection occurring during the first year of life is rather common among animals and plants, this source of bias may be common in studies estimating evolutionary parameters in wild populations (Hadfield, 2008; Nakagawa & Freckleton, 2008). The current study relies heavily on observing phenotypes that would have been expressed of individuals that had died: post-mortems of dead ewe lambs allow us to determine what their fecundity would have been the following spring. These data were instrumental for identifying the density-dependent manner in which pregnancy mediates total selection of body mass. This direct observation of a phenotype that would otherwise be missing is essentially a type of solution to the paradox of stasis, albeit one that may not be widely applicable in other systems.

Soay sheep have existed on an island, Soay, which is adjacent to Hirta, where the long-term study of Soay sheep occurs, since prehistoric times (Clutton-Brock & Pemberton, 2004a). Soay and Hirta have similar population dynamics, and as such, the associations among density, body size, pregnancy, and fitness reported here may be broadly representative of the long-term pattern selection experienced by Soay sheep. However, the current distribution of phenotype in the Soay sheep on Hirta, particularly their propensity to become pregnant in the first year, may be an adaptation to recent selection. Soay sheep were reintroduced to Hirta in 1932, when 107 individuals were moved from Soay (Clutton-Brock & Pemberton, 2004a), following the evacuation of the last human residents of Hirta. During the subsequent decades of much lower population size (Clutton-Brock et al., 2004), selection likely favoured a much faster life history. While results presented here are derived from data for the range of observed population sizes since 1985, they are strongly indicative of what selection would happen at lower population densities. At lower population densities, survival rates of ewe lambs would be very high, regardless of pregnancy status (Figure A1). Additionally, the survival of offspring of ewes that became pregnant as lambs is extremely strongly density dependent (Table B5), and as such there would be a clear benefit, and little or no survival cost, to early reproduction at smaller population sizes, such as those experienced by Soay sheep on Hirta during a key period in the last century.

## Summary

This study provides evidence of a trade-off constraining the evolution of body mass in an unmanaged population of an ungulate species. The Soay sheep population of Hirta has been established as an example of the paradox of stasis, and our results partly elucidate why selection of body mass may not be as strong as it initially appears. We show that selection towards larger body sizes would be stronger if not for the positive genetic correlation between lamb body mass and early pregnancy, and their opposing associations with fitness. Large lamb body mass in females is associated with higher rates of first-year survival and lifetime rearing success, but also with a higher chance of early pregnancy. The latter has a very large viability cost unless population density is low. More generally, this study underscores how a mechanistic understanding of life history patterns allows for a more fine-grained understanding of natural selection, with the potential to explain evolutionary conundrums such as the paradox of stasis.

## Acknowledgements

The authors first wish to thank all project members and many volunteers for their contributions to field work on St. Kilda, and to all those who have contributed to keeping the project running over many years, including T. Clutton-Brock, M. Crawley, S. Albon, T. Coulson, L. Kruuk, and D. Nussey. The authors are also grateful to the National Trust for Scotland and Scottish Natural Heritage for permission to work on St. Kilda, and QinetiQ and Eurest for logistics and other support on the island. M. J. Janeiro was supported by a PhD scholarship (SFRH/BD/96078/2013) funded by the Fundação para a Ciencia e Tecnologia (FCT). MBM is supported by a University Research Fellowship from the Royal Society (London).

## A Supplementary figure

**Figure A1:**
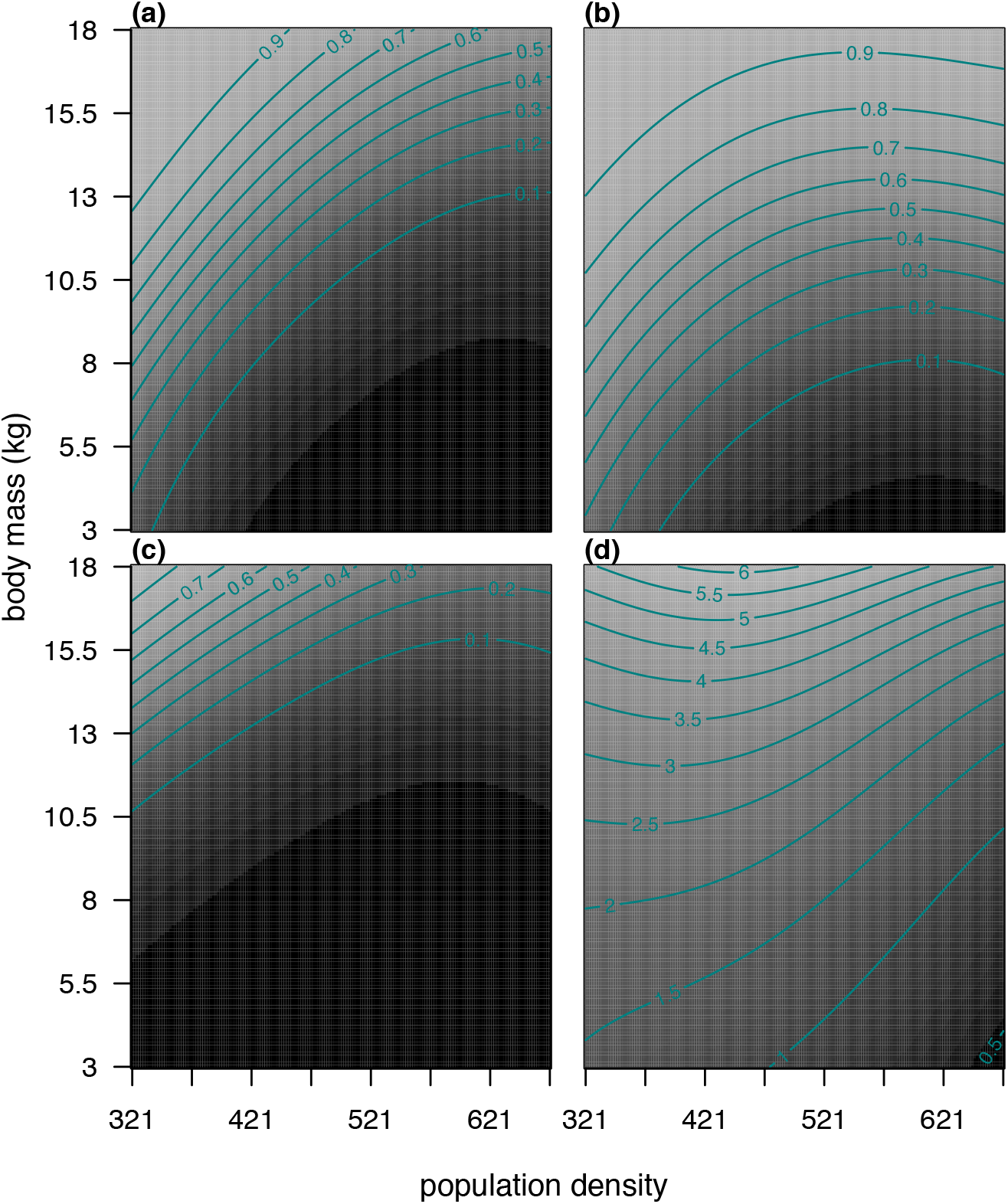
Probability of first-year survival in pregnant (a) and non-pregnant (b) ewe lambs, and AReS (c) and LReS (d) in Soay sheep ewes that survived their first annual cycle as a function of lamb body mass and population size. These are predictions based on models in Tables B4, B5 and B6.

## B Model summary tables

Lamb body mass in August is dependent on birth and measurement dates. Both are indicators of how long lambs had the opportunity to grow and are particularly relevant because higher growth rates occur during the warmer months (before August), when vegetation is most available (Crawley et al., 2004). As a result, Julian birth and measurement dates were included as covariates when modelling lamb body mass or lamb body mass was used as a predictor. Fixed effects for twinning status, population density and maternal age at parity, including quadratic terms for the latter two, were included in all regressions, as well as the following linear interaction terms: between lamb body mass and population density in models including lamb body mass as predictor, between early pregnancy status and population density in models including early pregnancy as a predictor, and between these three traits in models including both lamb body mass and early pregnancy status as predictors. Likewise, random effects to estimate among-mother and among-cohort variation were included in all regressions. This fixed and random structure was applied to all statistical models except the one modelling AReS (due to a lack of statistical power).

**Table B1:** Coefficients of a linear regression of August body mass in Soay ewe lambs. All covariates, except for twin status, were mean-centred. Variances in breeding values, random cohorts and maternal identity were estimated. 95% credible intervals correspond to HPD intervals. See Eq. (1) (main text) for model description.

| parameter | posterior mode | 95% CrI |
| --- | --- | --- |
| intercept | 1.60 | (1.02; 2.07) |
| density ($\times 100$) | -0.27 | (-0.59; 0.05) |
| density<sup>2</sup> ($\times 100$) | 0.00 | (-0.01; 0.00) |
| twin | -2.91 | (-3.21; -2.62) |
| maternal age | 0.36 | (0.29; 0.40) |
| maternal age<sup>2</sup> | -0.11 | (-0.13; -0.10) |
| birth date | -0.06 | (-0.08; -0.05) |
| measurement date | 0.09 | (0.04; 0.13) |
| additive genetic variance: | 0.67 (0.07; 1.14) |  |
| variance in cohort effects: | 0.71 (0.33; 1.08) |  |
| variance in maternal effects: | 0.59 (0.29; 1.21) |  |
| residual variance: | 1.81 (1.47; 2.33) |  |

**Table B2:**
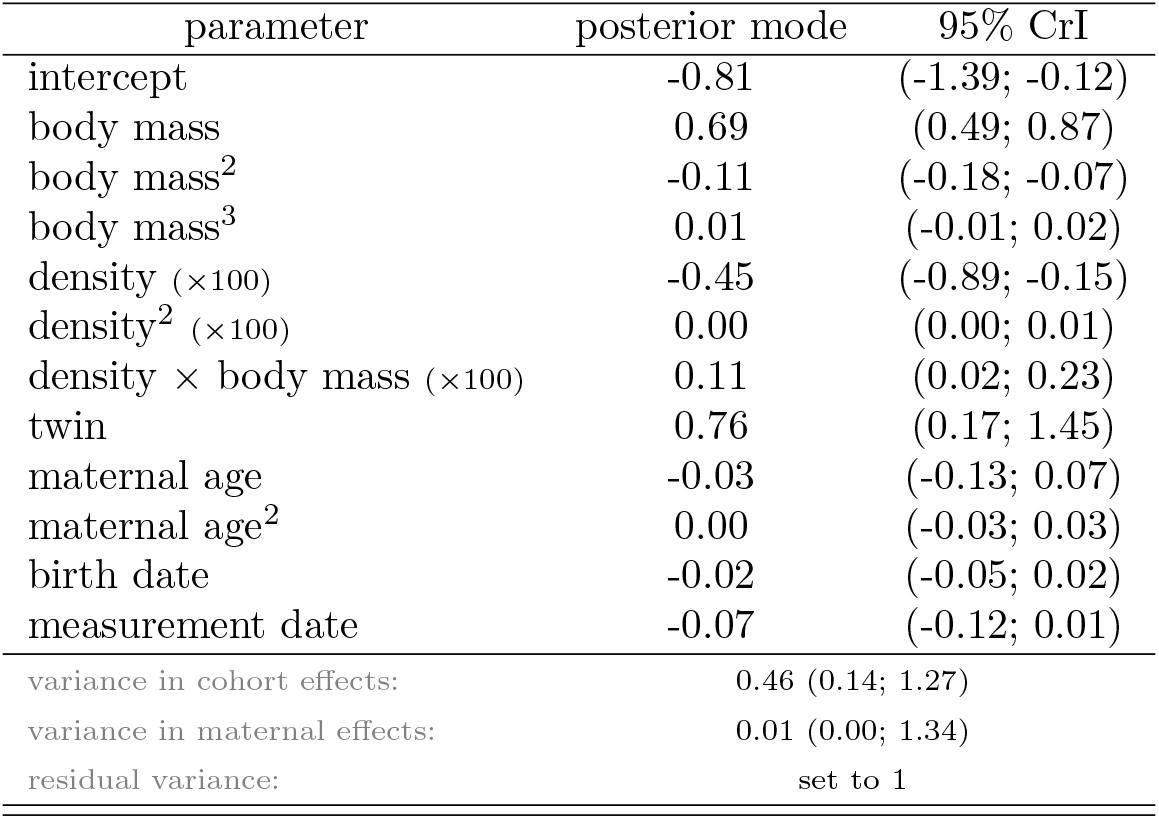
Coefficients of a binomial regression (logit link function) of early pregnancy on body mass in Soay ewe lambs. Other fixed effects include population density, twin status, and maternal age. All covariates, except for twin status, were mean-centred. Variances in random cohorts and maternal identity were also estimated. 95% credible intervals correspond to HPD intervals. See Eq. (2) (main text) for model description.

**Table B3:** Coefficients of the bivariate model of lamb body mass and early pregnancy in Soay sheep ewes. See Eq. (3) and (4) (main text) for model description.

| parameter |  | posterior mode | 95% CrI |
| --- | --- | --- | --- |
| fixed effects |  |  |  |
| body mass | intercept | 13.821 | (13.215; 14.261) |
|  | density | -0.002 | (-0.005; 0.001) |
|  | density<sup>2</sup> | 0.000 | (0.000; 0.000) |
|  | twinning | -2.972 | (-3.245; -2.652) |
|  | maternal age | 0.391 | (0.330; 0.436) |
|  | maternal age<sup>2</sup> | -0.108 | (-0.125; -0.095) |
| pregnancy | intercept | 0.000 | (-0.850; 0.636) |
|  | density | -0.005 | (-0.009; -0.001) |
|  | density<sup>2</sup> | 0.000 | (0.000; 0.000) |
|  | twinning | 1.851 | (1.055; 2.273) |
|  | maternal age | -0.140 | (-0.268; -0.023) |
|  | maternal age<sup>2</sup> | 0.043 | (-0.003; 0.071) |
| random effects |  |  |  |
| breeding values | body mass | 0.785 | (0.199; 1.272) |
|  | body mass:pregnancy | 0.307 | (-0.001; 1.494) |
|  | pregnancy | 0.695 | (0.001; 5.859) |
| cohort | body mass | 0.596 | (0.266; 1.141) |
|  | body mass:pregnancy | 0.385 | (0.136; 1.163) |
|  | pregnancy | 0.932 | (0.136; 1.163) |
| maternal ID | body mass | 0.577 | (0.349; 1.014) |
|  | body mass:pregnancy | 0.002 | (-0.159; 0.414) |
|  | pregnancy | 0.005 | (0.000; 0.824) |
| residuals | body mass | 1.940 | (1.387; 2.224) |
|  | body mass:pregnancy | 1.059 | (0.699; 1.267) |
|  | pregnancy | 1.000 | (1.000; 1.000) |

**Table B4:** Coefficients of mixed effect binomial regressions with logit link function exploring the association of first-year survival with size and pregnancy in Soay ewes; (a) size-dependent first-year survival, (b) first-year survival as a function of size and pregnancy. Covariates except for twin and pregnancy status were mean-centred. See Eq. (10) (main text) for model description.

| parameter | posterior mode | 95% CrI |
| --- | --- | --- |
| (a) |  |  |
| intercept | -0.55 | (-1.59; 0.38) |
| body mass | 0.50 | (0.36; 0.66) |
| density ($\times 100$) | -1.38 | (-1.95; -0.72) |
| density<sup>2</sup> ($\times 100$) | 0.01 | (0.00; 0.01) |
| density × body mass ($\times 100$) | -0.06 | (-0.18; 0.05) |
| twin | 0.06 | (-0.70; 0.74) |
| maternal age | -0.02 | (-0.12; 0.12) |
| maternal age<sup>2</sup> | 0.01 | (-0.03; 0.05) |
| birth date | 0.02 | (-0.02; 0.05) |
| measurement date | -0.10 | (-0.20; -0.03) |
| variance in cohort effects: 0.78 (0.78; 4.18) |  |  |
| variance in maternal effects: 1.82 (0.61; 4.22) |  |  |
| residual variance: set to 1 |  |  |
| (b) |  |  |
| intercept | 0.14 | (-0.95; 1.23) |
| pregnancy | -2.15 | (-2.73; -1.35) |
| body mass | 0.69 | (0.47; 0.89) |
| body mass × pregnancy | 0.19 | (-0.18; 0.43) |
| density ($\times 100$) | -0.89 | (-1.75; -0.28) |
| density<sup>2</sup> ($\times 100$) | 0.00 | (0.00; 0.01) |
| density × pregnancy ($\times 100$) | -1.35 | (-2.04; -0.70) |
| density × body mass ($\times 100$) | 0.10 | (-0.07; 0.24) |
| density × body mass × pregnancy ($\times 100$) | -0.04 | (-0.34; 0.27) |
| twin | 0.19 | (-0.74; 0.88) |
| maternal age | 0.04 | (-0.12; 0.14) |
| maternal age<sup>2</sup> | 0.00 | (-0.04; 0.04) |
| birth date | 0.01 | (-0.03; 0.05) |
| measurement date | -0.15 | (-0.23; -0.04) |
| variance in cohort effects: 2.04 (0.98; 5.11) |  |  |
| variance in maternal effects: 2.23 (0.85; 4.89) |  |  |
| residual variance: set to 1 |  |  |

**Table B5:** Coefficients of a binomial regression (logit link function) of AReS in surviving ewe lambs on lamb body mass, birth and measurement dates, and population density in Soay sheep ewe lambs. All covariates were mean-centred and the 95% credible intervals correspond to HPD intervals. See Eq. (11) (main text) for model description.

| Parameter | Posterior mode | 95% CrI |
| --- | --- | --- |
| intercept | -4.00 | (-5.08; -2.94) |
| body mass | 0.64 | (0.43; 1.02) |
| density ($\times 100$) | -1.06 | (-2.41; -0.11) |
| density<sup>2</sup> ($\times 100$) | 0.00 | (0.00; 0.01) |
| birth date | 0.03 | (-0.04; 0.10) |
| measurement date | 0.06 | (-0.11; 0.16) |
| residual variance: set to 1 |  |  |

**Table B6:** Coefficients of Poisson regressions with logarithm link functions, exploring the association between LReS in ewes surviving their first year and lamb body mass and early pregnancy; (a) size-dependent LReS, and (b) LReS as a function of lamb August body mass and early pregnancy. Covariates except for twin status and early pregnancy status were mean-centred, and 95% credible intervals correspond to HPD intervals. See Eq. (12) (main text) for model description.

| parameter | posterior mode | 95% CrI |
| --- | --- | --- |
| (a) |  |  |
| intercept | 0.64 | (0.12; 0.99) |
| body mass | 0.14 | (0.04; 0.23) |
| density ($\times 100$) | -0.15 | (-0.49; 0.01) |
| density<sup>2</sup> ($\times 100$) | 0.00 | (0.00; 0.00) |
| density × body mass ($\times 100$) | 0.04 | (-0.06; 0.09) |
| twin | 0.07 | (-0.38; 0.52) |
| maternal age | 0.00 | (-0.08; 0.06) |
| maternal age<sup>2</sup> | 0.01 | (0.00; 0.03) |
| birth date | 0.01 | (-0.02; 0.03) |
| measurement date | 0.01 | (-0.06; 0.07) |
| variance in cohort effects: | 0.00 (0.00; 0.37) |  |
| variance in maternal effects: | 0.38 (0.00; 0.68) |  |
| residual variance: | 0.45 (0.20; 0.86) |  |
| (b) |  |  |
| intercept | 0.38 | (-0.07; 0.83) |
| pregnancy | 0.37 | (-0.06; 0.82) |
| body mass | 0.14 | (0.03; 0.27) |
| pregnancy × body mass | 0.02 | (-0.28; 0.09) |
| density ($\times 100$) | -0.25 | (-0.52; 0.06) |
| density<sup>2</sup> ($\times 100$) | 0.00 | (0.00; 0.00) |
| density × pregnancy ($\times 100$) | 0.34 | (-0.09; 0.75) |
| density × body mass ($\times 100$) | 0.10 | (0.00; 0.21) |
| density × body mass × pregnancy ($\times 100$) | -0.24 | (-0.39; -0.06) |
| twin | 0.14 | (-0.34; 0.57) |
| maternal age | 0.00 | (-0.08; 0.07) |
| maternal age<sup>2</sup> | 0.02 | (0.00; 0.04) |
| birth date | 0.00 | (-0.02; 0.03) |
| measurement date | 0.02 | (-0.05; 0.07) |
| variance in cohort effects: | 0.00 (0.00; 0.38) |  |
| variance in maternal effects: | 0.30 (0.00; 0.60) |  |
| residual variance: | 0.51 (0.26; 0.91) |  |

**Table B7:**
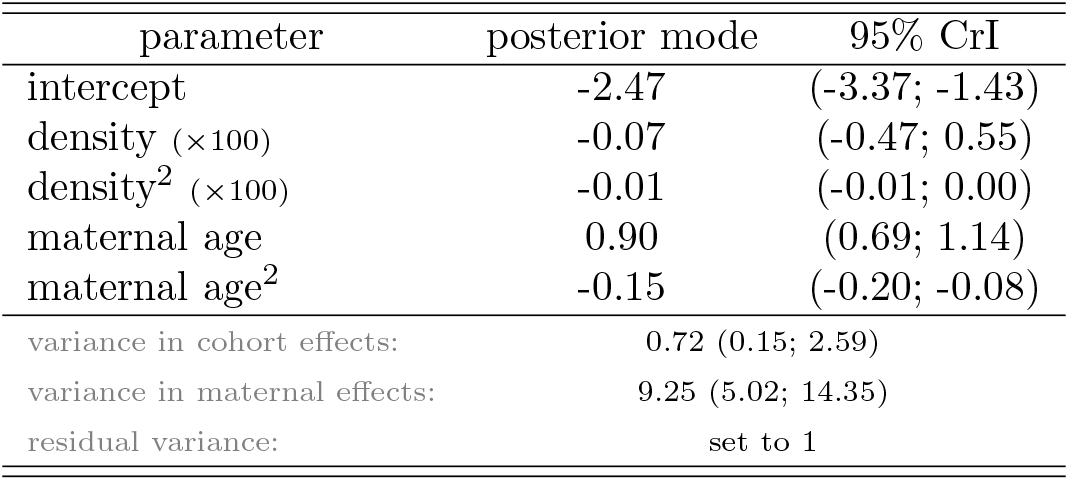
Coefficients of a binomial regression (logit link function) of the probability of twinning as a function of mean-centred population density and maternal age. 95% credible intervals correspond to HPD intervals. See Eq. (13) (main text) for model description.

**Table B8:** Coefficients of a random regression animal model to estimate additive genetic reaction norms of early pregnancy on body mass in Soay ewe lambs. A binomial regression with logit link function was fit to the data. All covariates were mean-centred. Variance in random cohorts was also estimated. 95% credible intervals correspond to HPD intervals.

| parameter | posterior mode | 95% CrI |
| --- | --- | --- |
| intercept | -0.88 | (-1.63; -0.17) |
| body mass | 0.74 | (0.53; 0.98) |
| density ($\times 100$) | -0.46 | (-1.02; -0.08) |
| density<sup>2</sup> ($\times 100$) | 0.00 | (0.00; 0.01) |
| variance in breeding values (intercepts): | 2.04 (0.13; 9.19) |  |
| variance in breeding values (slopes for mass): | 0.19 (0.05; 0.67) |  |
| covariance between breeding values (intercepts and slopes): | 0.61 (0.15; 2.35) |  |
| variance in cohort effects: | 0.80 (0.19; 2.25) |  |
| residual variance: | set to 1 |  |

## C Details on solving derivatives and integrals

Estimating average trait values (Eq. (5), main text) and variances (Eq. (6), main text) on the expected scale implies solving integrals of high dimensionality in the context of the path analysis, and particularly when these equations are applied to the vector-valued function ***f*_(*l*)_** in equation (14). In such circumstances, solving integrals using Monte Carlo simulations is particularly useful as the error is independent from the integrand dimensionality, but mainly because other numerical methods are likely to fail. Monte Carlo integration relies on the fact that the integral ∫*_L_ g*(*l*)*f*(*l*)*dl* (or ∫*_L_ g*^−1^(*l*)*f*(*l*)*dl* if applied to an inverse link function) corresponds to the expectation of *g*, 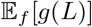. As a consequence, a random sample of latent values, *L*, generated using its density *f* can be used to approximate the empirical average of *g*,

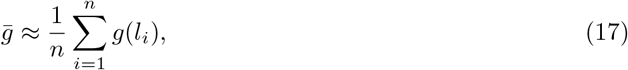

where *n* is the number of draws from *L*. 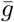 is also an approximation to ∫*_L_ g*(*l*)*f*(*l*)*dl*. This approximation converges to the real value of the integral by the law of large numbers as *n* → ∞ (Robert & Casella, 2005). We sampled a large enough number of values from the distribution of the latent traits on which we evaluated ***f*(*l***). Since the statistical models were fitted in a Bayesian framework, this procedure was applied to *n_post_* = 2000 posterior independent samples.

For lower dimensional integrals, and particularly to obtain Φ (average derivative of the expected values with respect to the latent ones) in equation (8) (main text), we used function *cuhre* from R package **R2Cuba** that implemments a deterministic algorithm that uses cubature rules of polynomial degree, also allowing for multidimensional integration.

Likewise, the differentiation in the calculation of extended selection gradients (Eq. 15, main text) was accomplished by numerical linear approximation,

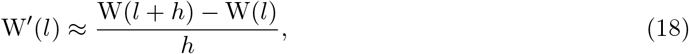

where *h* is some small value. So, the extended selection gradient for mass would be obtained by evaluating W for a certain value of latent mass *l* and at for *l* + *h*. This procedure was applied to *n_MC_* = 2000 values from the distribution of the latent distributions on which we evaluated ***f*(*l***) times *n_post_* = 2000 posterior independent samples.

